# The long non-coding RNA *LINDA* restrains cellular collapse following DNA damage in *Arabidopsis thaliana*

**DOI:** 10.1101/2023.06.28.546876

**Authors:** Josephine Herbst, Solveig Henriette Nagy, Ilse Vercauteren, Lieven De Veylder, Reinhard Kunze

**Author notes:** **ORCID IDs:** 0000-0002-6253-4927 (J.H.); 0000-0001-9561-8468 (I.V.); 0000-0003-1150-4426 (L.D.V.); 0000-0002-3304-5550 (R.K.).

## Abstract

The genomic integrity of every organism is endangered by various intrinsic and extrinsic stresses. To maintain the genomic integrity, a sophisticated DNA damage response (DDR) network is activated rapidly after DNA damage. Notably, the fundamental DDR mechanisms are conserved in eukaryotes. However, knowledge about many regulatory aspects of the plant DDR is still limited. Important, yet little understood, regulatory factors of the DDR are the long non-coding RNAs (lncRNAs). In humans, 13 lncRNAs functioning in DDR have been characterized to date, whereas no such lncRNAs have been characterized in plants yet. By meta-analysis, we identified the *long intergenic <u>n</u>on-coding RNA induced by DNA <u>da</u>mage* (*LINDA*) that responds strongly to various DNA double-strand break-inducing treatments, but not to replication stress induced by mitomycin C. After DNA damage, *LINDA* is rapidly induced in an ATM- and SOG1-dependent manner. Intriguingly, the transcriptional response of *LINDA* to DNA damage is similar to that of its flanking hypothetical protein-encoding gene. Phylogenetic analysis of putative Brassicales and Malvales *LINDA* homologs indicates that *LINDA* lncRNAs originate from duplication of a flanking small protein-encoding gene followed by pseudogenization. We demonstrate that *LINDA* is not only needed for the regulation of this flanking gene, but also for fine-tuning of the DDR after the occurrence of DNA double-strand breaks. Moreover, Δ*linda* mutant root stem cells are unable to recover from DNA damage, most likely due to hyper-induced cell death.

**SIGNIFICANT STATEMENT:** We unraveled the functional relevance of the first lncRNA within the DNA damage response network of *Arabidopsis thaliana.* This lncRNA, termed *LINDA*, is an important part of the DNA damage response network, as it is needed for accurate regulation of cell death and cell cycle progression.

## INTRODUCTION

The ability to rapidly recognize and repair damaged DNA is critical for every living organism to maintain genome integrity. Common threats endangering genome integrity are extrinsic factors such as UV radiation and heavy metal pollution, and intrinsic factors such as reactive oxygen species (ROS). Consequently, a highly sophisticated signaling network, termed the DNA damage response (DDR) pathway, has evolved in all organisms. Notably, many factors of the DDR are conserved in eukaryotes, especially those that are involved in recognition and signaling of the damage (Yoshiyama *et al*., 2013b). The key factors initiating the DDR are the two phosphoinositide 3-kinase-like kinases ataxia telangiectasia-mutated (ATM) and ataxia telangiectasia and Rad3-related (ATR) that are both rapidly activated after DNA damage (Culligan *et al*., 2006). While ATM is activated by the MRN [MRE11 (meiotic recombination 11), RAD50 (radiation sensitive 50), NBS1 (Nijimegen breakage syndrome 1)] complex after DNA double-strand breaks (DSBs) (Lee and Paull, 2005, Culligan *et al*., 2006), ATR is predominantly activated by single-strand breaks and replication stress (Culligan *et al*., 2006). Once activated, both kinases can phosphorylate the central integrator of the DDR, thus inducing transcriptional changes (Yoshiyama *et al*., 2013a, Ogita *et al*., 2018). In humans, this integrator is the tumor suppressor protein p53, while the non-related transcription factor SUPPRESSOR OF GAMMA IRRADIATION 1 (SOG1) fulfills this task in plants (Yoshiyama *et al*., 2009). During the DDR, SOG1 gets hyperphosphorylated and binds directly to a conserved motif within the promoter of at least 300 target genes in *Arabidopsis thaliana* (Yoshiyama *et al*., 2013a, Yoshiyama *et al*., 2017). The consensus sequence of the *cis*-acting SOG1-binding motif is a palindromic CTT[N_7_]AAG sequence that was identified by chromatin immunoprecipitation sequencing (ChIP-Seq) analysis. This motif is enriched in the −400 bp promoter regions of a large fraction of SOG1-dependent early DDR genes (Bourbousse *et al*., 2018, Ogita *et al*., 2018). SOG1 target genes are predominantly involved in DNA repair, cell cycle checkpoint activation, and signal transduction. For instance, SOG1 activates different cyclin-dependent kinase inhibitors, thus inhibiting cell cycle progression (Yoshiyama *et al*., 2009, Weimer *et al*., 2016, Chen *et al*., 2017, Ogita *et al*., 2018). Moreover, hyperphosphorylated SOG1 activates *BREAST CANCER SUSCEPTIBILITY GENE 1* (*BRCA1*) and *RADIATION SENSITIVE 51* (*RAD51*), both involved in DNA repair via homologous recombination (Yoshiyama *et al*., 2017, Ogita *et al*., 2018). Besides the activation of genes involved in DNA repair, one important aspect of the DDR is the selective activation of programmed cell death, for example in stem cells of the root meristem surrounding the quiescence center (Fulcher and Sablowski, 2009). Recent data showed the involvement of auxin and cytokinin in the activation of cell death triggered by DNA damage (Takahashi *et al*., 2021). Still, not much information about the molecular processes needed for the execution of cell death is available.

Although our knowledge of the plant-specific DDR has been tremendously expanded since the identification of SOG1, many questions regarding the fine-tuning of the DDR in response to various stresses remain unsolved. One group of biomolecules that are hypothesized to play an important role in this fine-tuning are non-coding RNAs. Non-coding RNAs lack an apparent coding potential, exhibit low transcription rates and a high transcriptional turnover (Chen and Zhu, 2022). They are typically divided into two classes based on length: short non-coding RNAs (< 200 nt) and long non-coding RNAs (lncRNAs, > 200 nt) (Kapranov *et al*., 2007, Kapusta and Feschotte, 2014). Especially the lncRNAs have gained attention over the past years, because of their highly stress-specific transcriptional response (Di *et al*., 2014). Based on the so far analyzed lncRNAs in mammals and plants, it was shown that lncRNAs are frequently involved in the regulation of their target genes in *cis* or *trans* (Kopp and Mendell, 2018, Rai *et al*., 2019, Lucero *et al*., 2021). One of the most renowned lncRNAs in mammals is *Xist*, the mayor driver of X-chromosome inactivation (Brockdorff *et al*., 1991). Two well-studied plant lncRNAs from Arabidopsis are *COLD ASSISTED INTRONIC NONCODING RNA* (*COLDAIR*) and *COLD INDUCED LONG ANTISENSE INTRAGENIC RNA* (*COOLAIR*), both regulating the FLOWERING LOCUS C of Arabidopsis in response to cold stress via epigenetic modifications (Swiezewski *et al*., 2009, Kim and Sung, 2017). Another example in Arabidopsis and rice is *INDUCED BY PHOSPHATE STARVATION1* (*IPS1*). *IPS1* sequesters miRNA399 by target mimicry, thus releasing expression of the miRNA399 target gene *PHOSPHATE TRANSPORTER 2* (*PHO2*) in response to phosphate starvation (Franco-Zorrilla *et al*., 2007, Jabnoune *et al*., 2013).

So far, 13 human lncRNAs were shown to be involved in the transcriptional regulation of the DDR, the mediation of DNA-DNA or DNA-protein complexes assisting the binding and repair of DSBs, and the regulation of autophagy (Li *et al*., 2021). *Transcribed in the Opposite Direction of RAD51* (*TORAD*), for example, promotes the RAD51-dependent repair of DSBs (Gazy *et al*., 2015), while the *DNA Damage-Sensitive RNA1* (*DDSR1*) interacts with BRCA1 and modulates the DNA repair process (Sharma *et al*., 2015). In contrast to humans, no lncRNA functioning in the plant DDR was characterized yet. The first identification of lncRNAs involved in plant DDR was reported by Wang *et al*. in 2016. Here, 86 lncRNAs were found to be significantly up-or downregulated 3 h after treatment of *A. thaliana* with a sublethal dose of X-rays (Wang *et al*., 2016). Strikingly, the transcriptional activation for over 90% (79/86) of the identified lncRNAs depends on ATM, indicating the potential of plant lncRNAs to be important for the DDR to a similar extent as in humans. However, these lncRNAs were not characterized any further.

In this study, we performed a comparative meta-analysis of DNA damage-responsive lncRNAs in different published bulk RNA-Seq datasets. We found the lncRNA *At3g00800* to be strongly upregulated after induction of DSBs and termed it *<u>L</u>ong Intergenic <u>n</u>cRNA Induced by DNA <u>Da</u>mage* (*LINDA*). We demonstrated that *LINDA* is involved in the accurate execution of the DDR in *A. thaliana*. Moreover, we identified the putative target gene of *LINDA*, which is located downstream of the lncRNA-encoding gene.

## RESULTS

### Identification of lncRNAs regulated by DNA damage

To identify lncRNAs that are potentially involved in the DDR of *A. thaliana,* we performed a meta-analysis of different online available datasets (Wang *et al*., 2016, Huang *et al*., 2019, Czarnocka *et al*., 2020, Lee *et al*., 2021). We focused on DNA damage induced by either X-ray or UV-C.(Wang *et al*., 2016, Czarnocka *et al*., 2020) To exclude genes that might be differentially expressed due to high radiation or heat, we further included datasets for high light and mild heat stress (Huang *et al*., 2019, Lee *et al*., 2021). First, we analyzed all significantly (p < 0.05) up-or downregulated transcripts (Figure 1a) to ensure that no dataset was over-or underrepresented in our meta-analysis. Next, we analyzed all significantly (p < 0.05) up-or downregulated lncRNAs (Figure 1b). As described before (Di *et al*., 2014), the fraction of identified lncRNAs that are responsive to only one treatment was significantly higher in comparison to that of all transcripts (Figure S1). Intriguingly, only six lncRNAs were affected both by X-ray and UV-C (Figure 1b, red circle), of which the novel translated region *At3g00800* was the only one that is upregulated by both stresses (Figure 1c). Moreover, *At3g00800* was induced by the radiomimetic drugs bleomycin and zeocin, which trigger DSBs similar as X-rays (Figure 1d). The induction of *At3g00800* was directly proportional to the concentration of the radiomimetic drug (Figure 1e). However, mitomycin C, which primarily induces replication stress, did not affect *At3g00800* expression (Figure 1d). Moreover, *At3g00800* is induced by UV-C, but not UV-A or UV-B (Figure 1f), which primarily lead to the formation of pyrimidine dimers or DNA cross-links.

**Figure 1.**
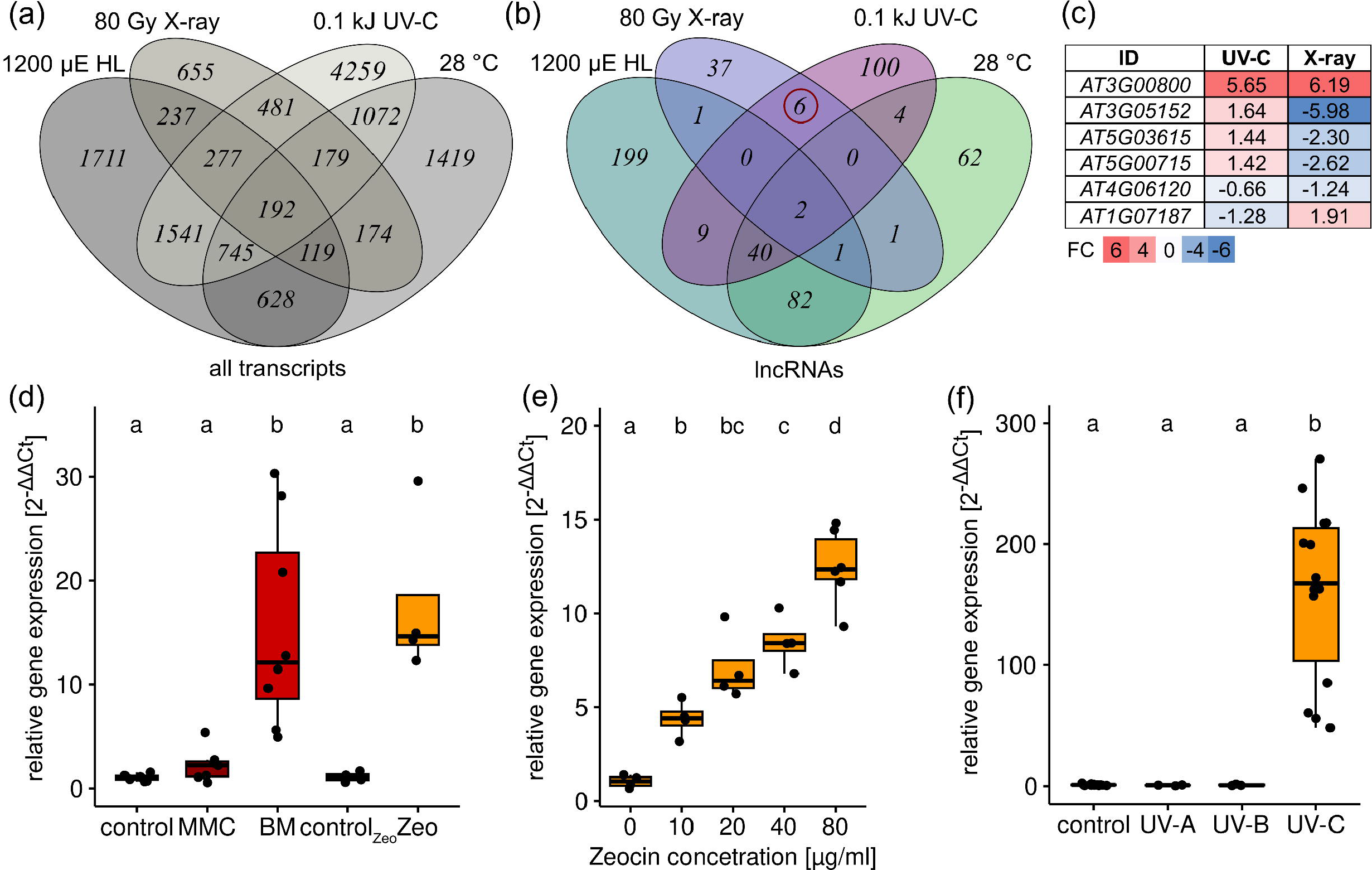
Comparative analysis of DNA damage-induced *Arabidopsis thaliana* transcriptomes. (a-b) Four RNA-seq datasets were analyzed for genes significantly (p < 0.05) regulated after DNA damage (by treatment with 1200 µE high light (HL), 80 Gy X-ray, 0.1 kJ UV-C or 28°C heat) with the Bioconductor DESeq2 package. The number of uniquely and commonly regulated genes were calculated and visualized by the CRAN VennDiagram package (a) for all transcripts and (b) for those, which are classified as long non-coding RNAs (lncRNAs). (c) The fold change (FC) of the six genes that are responsive to both X-ray and UV-C. (d) *LINDA* gene expression after different DNA-damaging treatments in 21-day-old *A. thaliana* seedlings, grown under short-day conditions and harvested 3 h after induction of either DSBs by bleomycin (BM) and zeocin (Zeo), or DNA cross-links by mitomycin C (MMC). (e) Transcriptional response of *LINDA* to increasing doses of zeocin, 3 h after onset of treatment. (f) Transcriptional response of *LINDA* to 2 kJ UV-A, UV-B or UV-C. Statistical significance was evaluated by one-way ANOVA, Sidak post-hoc. Distinct letters indicate statistically significant differences (p< 0.05, n ≥ 4).

### *LINDA* (*At3g00800*) is an early DDR gene and coexpresses with its flanking protein-encoding gene

Due to its location in the intergenic region between two protein-encoding genes (Figure 2a), we named our candidate gene *<u>L</u>ong Intergenic <u>n</u>cRNA Induced by DNA <u>Da</u>mage* (*LINDA*). Because DDR genes can be clustered into different categories, based on their temporal response to the applied damage (Bourbousse *et al*., 2018), we measured the transcript levels of *LINDA* at different time points after induction of DNA damage by either UV-C or zeocin. *LINDA* was significantly induced 60 min after UV-C exposure and 90 min following zeocin treatment, and transcripts remained detectable at high levels for several hours after the treatments (Figure 2b, c). Accordingly, *LINDA* belongs to the class of early response genes after DNA damaging treatments.

**Figure 2.**
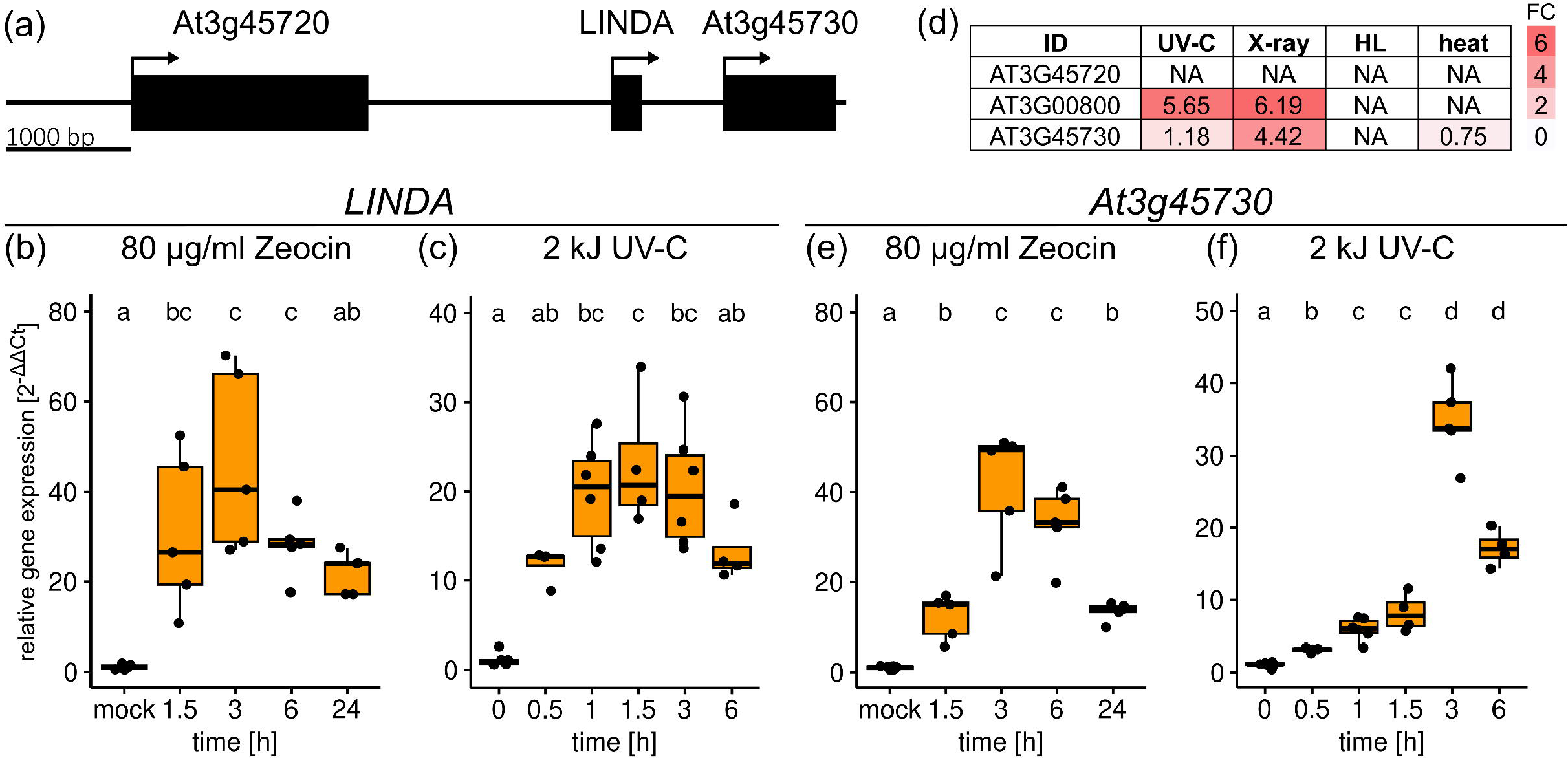
*LINDA* is an early response gene induced by various DNA-damaging treatments. (a) Schematic representation of the intergenic regions surrounding the *LINDA*-encoding gene. Black boxes indicate the annotated transcribed regions. (b-c) The gene expression in 21-day-old *A. thaliana* seedlings, grown under short-day conditions, analyzed after different DNA damaging treatments. Expression of *LINDA* was monitored at different time points during incubation with (b) 80 µg/ml zeocin or (c) after incubation with 2 kJ UV-C in comparison to mock treatments. (d) Fold change (FC) of *LINDA* (*At3g00800*) and the two flanking genes in the previously performed meta-analysis. (e-f) Transcriptional response of the hypothetical protein-encoding gene *At3g45730* at different time points after induction of DNA damage by treatment with 80 µg/ml zeocin (e) or 2 kJ UV-C (f). Statistical significance was evaluated by one-way ANOVA, Sidak post-hoc. Distinct letters indicate statistically significant differences (p< 0.05, n ≥ 4).

Because many lncRNAs regulate their flanking or overlapping genes (Ariel *et al*., 2015), we investigated the expression of the *LINDA* flanking genes in our initial meta-analysis. *LINDA* is located between a major facilitator superfamily protein-encoding gene (*At3g45720*) and a hypothetical protein-encoding gene (*At3g45730*) (Figure 2a). While the former was not affected by any of the applied stresses, *At3g45730* showed comparable expression patterns to *LINDA* after X-ray and UV-C treatment (Figure 2d) and was also an early response gene after zeocin treatment (Figure 2e). Although *At3g45730* was also significantly upregulated 30 min after UV-C treatment, the peak in gene expression was observed only 3 h after the treatment (Figure 2f). Thus, although *At3g45730* showed an altered accumulation kinetics after UV-C treatment in comparison to *LINDA* (Figure 2c, f), both genes were significantly induced at very early time points after DNA damage, suggesting a regulatory connection between both genes.

### *LINDA* is controlled by the ATM-SOG1 pathway

We analyzed the intergenic regions flanking *LINDA* and identified two SOG1-binding motifs within the putative *LINDA* promoter region (Figure 3a). Moreover, two additional SOG1-binding motifs were found in the 5’ untranslated region (5’ UTR) and the coding sequence (CDS) of *At3g45730* (Figure 3a). Prior ChIP-Seq experiments by Bourbousse *et al*. (2018), using a pSOG1:SOG1-3xFLAG construct as bait, had shown that SOG1 binds to the promoter region of *At3g00800* (*LINDA*) and the 5’ UTR of *At3g45730*. To verify that *LINDA* and *At3g45730* are indeed controlled by SOG1, we examined the transcript levels of both genes in *atm*, *atr*, and *sog1-1* mutants after either zeocin or UV-C treatment (Figure 3b-f). Three hours after zeocin treatment, both *LINDA* and *At3g45730* were strongly induced in wild-type and *atr* mutant plants, while no response in *sog1-1* and *atm* mutants was detectable (Figure 3b, c). These results demonstrate that both *LINDA* and *At3g45730* are controlled by the ATM-SOG1 pathway. Similar experiments were performed with UV-C irradiation. Here, the expression of both genes was decreased by about 50% in the *atm* and *sog1-1* mutants (Figure 3d, e). Thus, both genes are only partially dependent on the ATM-SOG1 pathway after UV-C treatment. At later time points after UV-C treatment, the transcript level of *At3g45730* recovered in the *sog1-1* mutant to the wild-type level, indicating that the response in this mutant is only delayed (Figure 3f). Notably, *LINDA* was hyperactivated in the *atr* mutant under both zeocin and UV-C treatments (Figure 3b, d), suggesting an ATR-dependent regulation of *LINDA*, independent of SOG1.

**Figure 3.**
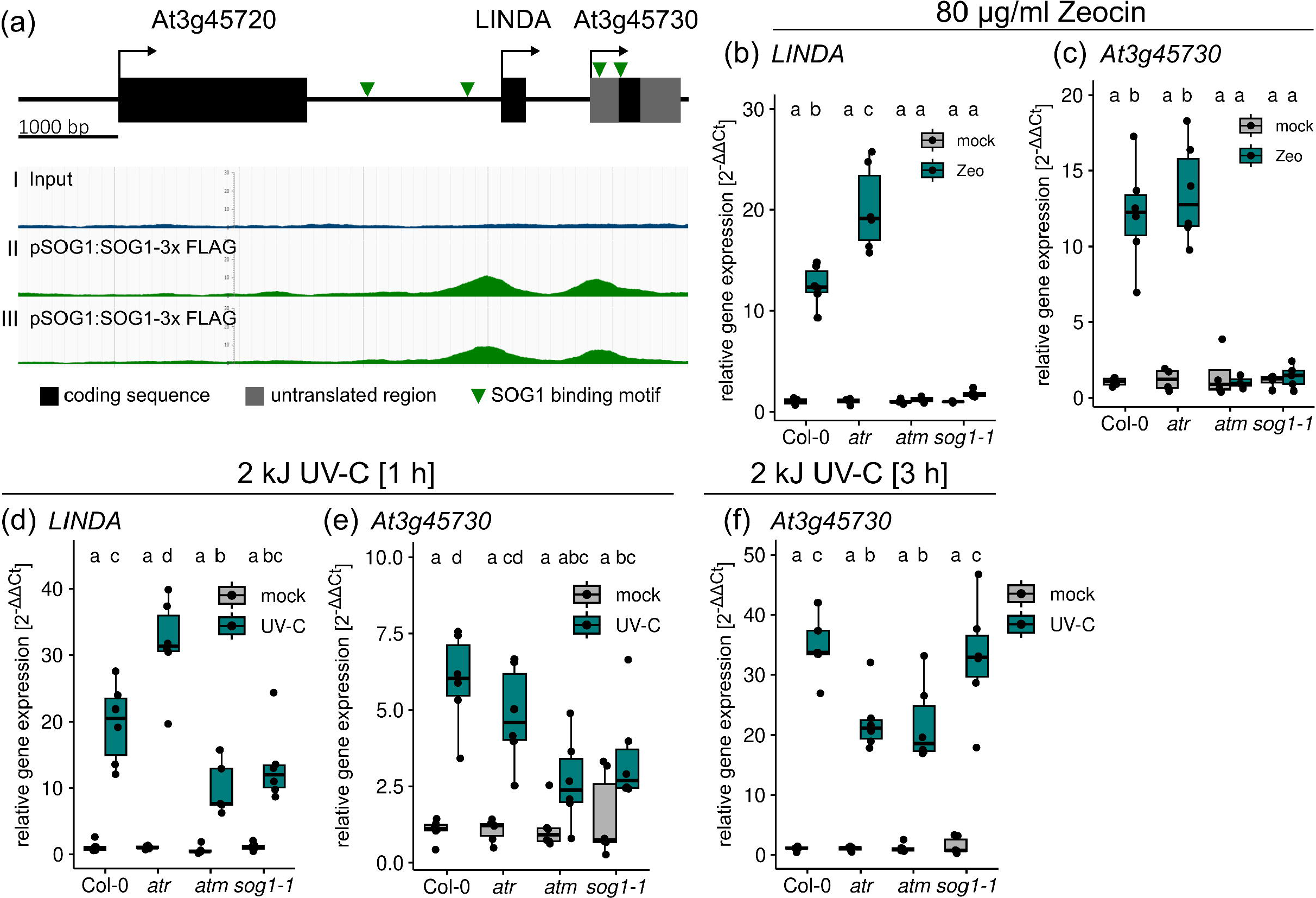
*LINDA* is directly regulated by the ATM-SOG1 pathway after induction of DSBs. (a) Schematic representation of the *LINDA* genome region. Two SOG1-binding motifs (green arrowheads) were identified in the putative promoter region of *LINDA*, as well as in the 5’ untranslated region (5’ UTR) and the coding sequence of the flanking hypothetical protein-encoding gene. ChiP-Seq data, available on the plantseq.org server, using pSOG1:SOG1-3xFLAG as bait showed that SOG1 can bind both *LINDA* and *At3g45730* (Bourbousse *et al*., 2018). Two replicates for ChiP-Seq data are depicted (II, III) in comparison to the used input without using pSOG1:SOG1-3xFLAG as bait (I). (b-c) Expression of *LINDA* (b) and *At3g45730* (c) in wild type (Col-0), *atr, atm* and *sog1-1* seedlings 3 h after treatment with 80 µg/ml zeocin. (d) Expression of *LINDA* 1 h after a 2-kJ UV-C treatment. (e-f) Expression of *At3g45730* in wild type, *atr, atm* and *sog1-1* seedlings 1 h (e) and 3 h (f) after a 2-kJ UV-C treatment. Statistical significance was evaluated by two-way ANOVA, Sidak post-hoc. Distinct letters indicate statistically significant differences (p< 0.05, n ≥ 4).

### *LINDA* presumably evolved by gene duplication of the flanking *At3g45730* gene

The gene *At3g45730* located downstream of *LINDA* encodes a yet uncharacterized hypothetical protein. As the transcription of this gene is also induced by various DNA-damaging stresses, like *LINDA*, we investigated the relationship between both genes. First, we searched for putative homologs of the hypothetical protein. A BLAST search using the *At3g45730* amino acid sequence as query uncovered the existence of several homologs in various Brassicales and Malvales species, e.g., *Brassica rapa* and *Gossypium hirsutum,* but not in Arabidopsis (Figure 4a, c). Sequence alignment of the putative homologs revealed a conserved PP[K/R]RG motif in the center of the protein sequence and a conserved serine residue at the C-terminus (Figure 4a). Using Alphafold, we visualized the putative protein structure, even though the confidence level of parts of the protein was rather low (Figure 4b). The two proline residues of the conserved PP[K/R]RG motif cause a kink in the helical structure, leading to the exposure of the downstream [K/R]RG residues (Figure 4b, inset). The conserved serine residue was found to be phosphorylated in *A. thaliana* (Roitinger *et al*., 2015). In summary, the sequence alignment pointed towards functionally relevant and therefore conserved features of the hypothetical protein.

**Figure 4.**
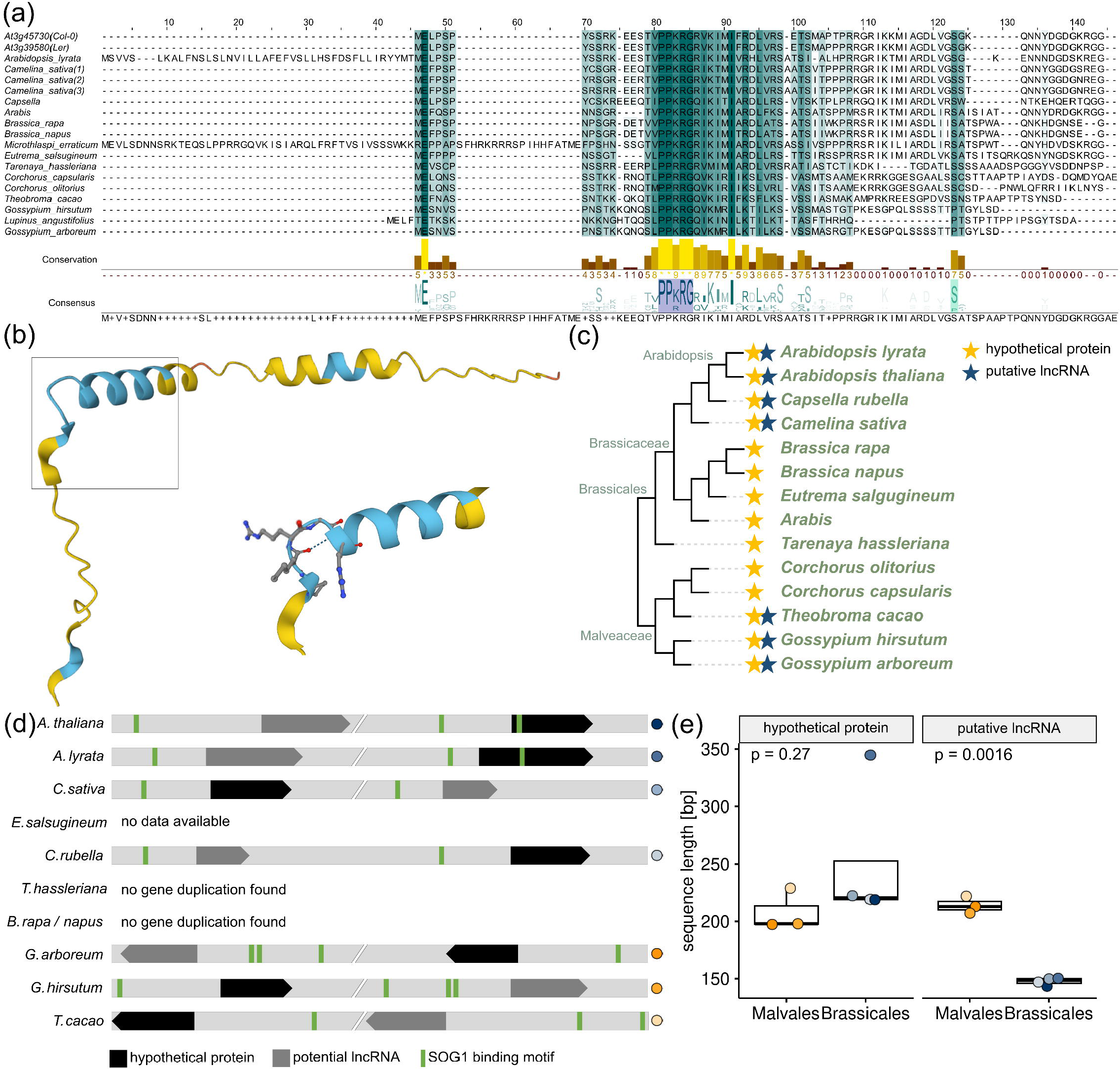
*In silico* analysis of the hypothetical At3g45730 protein. (a) Protein alignment of At3g45730 for *A. thaliana* ecotype Col-0 (top line) with homologs in *Arabidopsis lyrata* (v.1.0), *Arabidopsis thaliana* (TAIR10.1), *Brassica napus* (Bra_napus_v2.0), *Brassica rapa* (CAAS_Brap_v3.01), *Camelina sativa* (Cs), *Capsella rubella* (Caprub1_0), *Eutrema salsugineum* (Eutsalg1_0), *Gossypium arboreum* (Gossypium_arboreum_v1.0), *Gossypium hirsutum* (Gossypium_hirsutum_v2.1), *Tarenaya hassleriana* (ASM46358v1), *Theobroma cacao (*Criollo_cocoa_genome_V2). The conservation histogram is given below, depicting the highest degree of conservation in bright yellow. The consensus sequence of all putative homologs is given at the bottom. Conserved motifs are highlighted in dark and light blue. (b) Predicted protein structure by AlphaFold, with enlargement of the conserved PP[K/R]RG motif. Regions highlighted in blue correspond to a confident prediction with a per-residue confidence score (pLDDT) between 90 and 70, while yellow regions correspond to a low confidence (70 > pLDDT > 50). (c) Comparative phylogenetic analysis for the presence of *LINDA* (blue star) and the hypothetical At3g45730 protein (yellow star) in different Brassicales and Malvales species. (d) Schematic representation of the putative hypothetical protein homologs with their gene duplicate, putatively coding for a lncRNA similar to *LINDA*. (e) Lengths of the coding sequences of the hypothetical protein and the duplicated small open reading frame/putative lncRNA in Brassicales and Malvales species.

Next, we performed a BLASTN search using the transcribed *At3g45730* nucleotide sequence, which revealed that one of the closest homologs is the lncRNA *LINDA*. Indeed, *LINDA* contains a short open reading frame (ORF), which would code for a 48-amino acid-long protein with high similarity to the coding sequence of the hypothetical protein (Figure S2). We therefore hypothesize that *LINDA* originated from the duplication of the neighboring *At3g45730* gene, followed by pseudogenization. To strengthen this hypothesis, we screened the flanking regions of the putative *At3g45730* homologs, found in our earlier BLAST analysis, for duplicated sequences or annotated lncRNAs. Strikingly, we could identify putative gene duplications in the flanking intergenic region of 70% (7/10) of the investigated species, with available genome assembly data on the National Center for Biotechnology Information Genome Viewer server (Figure 4d). Moreover, the transcription direction and the presence of at least one SOG1-binding motif within the −1000 bp region of both the *At3g45730* homologous genes and the gene duplicate/lncRNAs, appear to be conserved (Figure 4d). Comparing the sequence length of the small ORF of the putative lncRNA and the CDS of the hypothetical protein between the different species, revealed that while the CDS of the hypothetical protein showed similar lengths, the lengths of the small ORF/putative lncRNA were shortened in Brassicales species (Figure 4e). The identification of lncRNAs in other non-model species is lacking behind *A. thaliana*. Therefore, we cannot confirm if those gene duplicates in other species do indeed encode lncRNAs. However, based on the phylogenetic analysis, it is likely that the gene duplication and pseudogenization did not only appear in Arabidopsis, but also other plant species.

### Transcriptome analysis analyses of the wild type and **Δ***linda* mutant response to DSBs

To examine the physiological function of *LINDA in planta*, we created a CRISPR/Cas12 deletion mutant using a dual-guide approach (Wolter and Puchta, 2019). The mutant, termed Δ*linda*, carries a 185-bp deletion, including 135 bp of the *LINDA* 3’ UTR (Figure 5a, b). Using deletion-spanning primers, we proved that the expression of *LINDA* is abolished in Δ*linda* mutant (Figure 5c).

**Figure 5.**
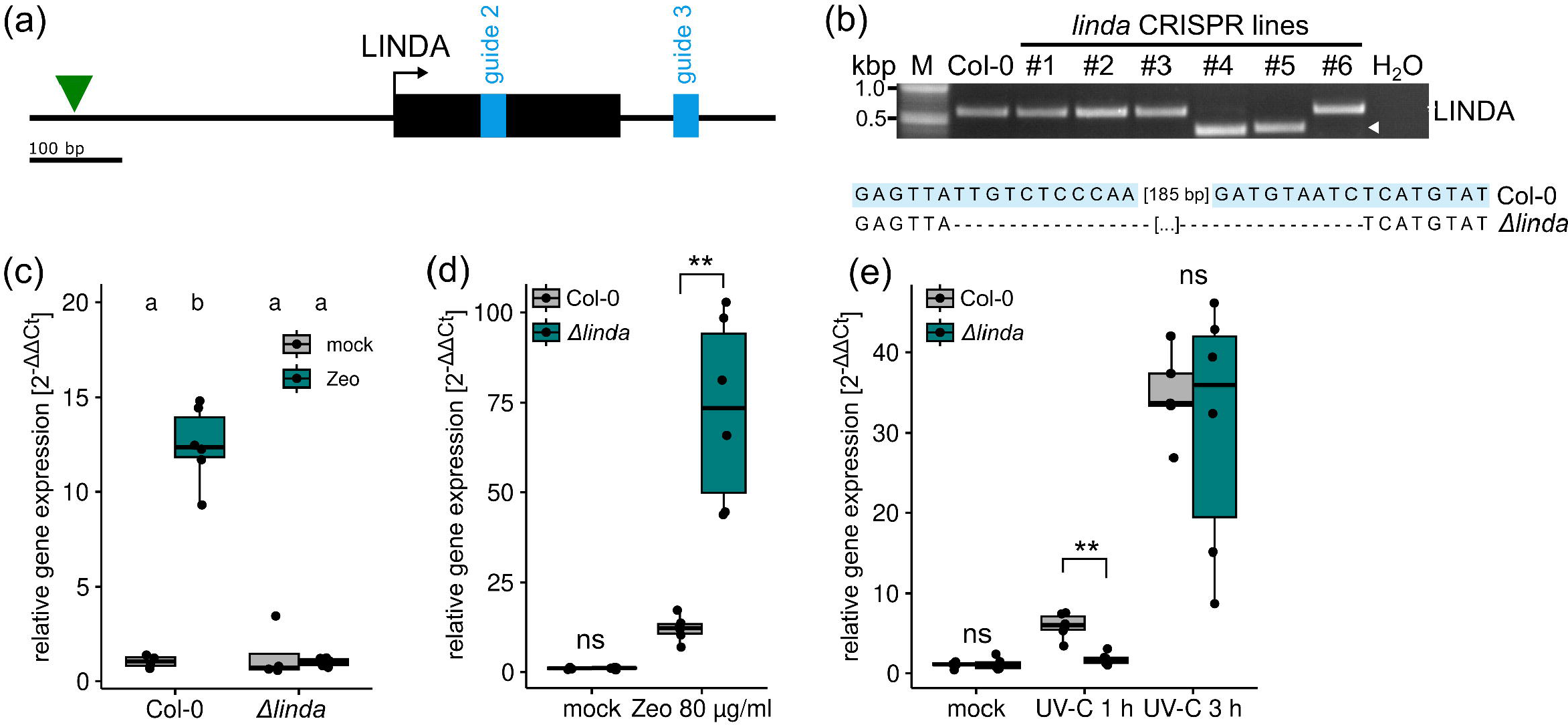
LINDA attenuates the expression of its flanking gene in response to DNA damage. (a) Schematic representation of the used guide RNAs (light blue) for construction of the Δ*linda* deletion mutant. The SOG1-binding site is depicted by the green triangle, while the transcribed region of *LINDA* is represented by the black box. (b-c) Verification of the *LINDA* deletion by PCR and sequencing (b), as well as by measuring the transcript levels using deletion spanning primers (c). Guide RNA sequences are highlighted in light blue. (d-e) Transcriptional activation of *At3g45730* in the Δ*linda* mutant after treatment with 80 µg/ml zeocin (d) or 2 kJ UV-C (e).

Initially, we analyzed the transcript levels of the flanking *At3g45730* gene in the Δ*linda* mutant after either UV-C or zeocin treatment. Intriguingly, after zeocin treatment, *At3g45730* transcripts were significantly (p = 0.0021) increased in the Δ*linda* mutant in comparison to the wild-type (Figure 5d). In contrast, after UV-C treatment, the expression of *At3g45730* was only reduced after 1 h in the Δ*linda* mutant and recovers to wild-type levels 2 h later (Figure 5e). Thus, *LINDA* might operate as a conditional repressor for its flanking gene.

Subsequently, to locate *LINDA* in the DDR regulatory network, we performed bulk RNA-Seq analyses of 21-day-old untreated and zeocin or UV-C treated wild-type and Δ*linda* plants. Principle component analyses (PCA) illustrate that the transcriptomes of untreated wild-type and Δ*linda* plants are quite similar (Figure S3a, b). Zeocin treatment leads not only to a severe reorganization of both wild-type and Δ*linda* transcriptomes (Figure S3a, PC1) but also reveals a significant difference between wild-type and mutant transcriptomes (Figure S3a, PC2). In contrast, UV-C treatment induces an even more pronounced reorganization of both wild-type and mutant transcriptomes (Figure S3b, PC1), but the stress-related transcriptomes of wild-type and Δ*linda* mutant plants are much more similar to each other than after zeocin treatment (Figure S3b, PC2). Thus, Δ*linda* might specifically impair the reaction to the DSB-inducing agent zeocin. This is further supported by the smaller number of commonly differentially regulated genes after zeocin treatment compared to UV-C treatment (Figure S3c, d). Although the absolute number of differentially regulated genes is higher after UV-C treatment, 70% of the identified responsive genes overlapped between Δ*linda* and the wild type (Figure S3d), whereas this is true for only 32% of the differentially regulated genes after zeocin treatment (Figure S3c). Moreover, the lack of *LINDA* has a stronger effect on the number of downregulated than that of the upregulated genes after zeocin treatment (Figure S3f). Most strikingly, the overlap of genes that are differentially regulated in the Δ*linda* mutant by both stresses is low (Figure S3e). This might indicate that *LINDA* is involved in the regulation of a different subset of target genes depending on the type of DNA damage.

Between the Δ*linda* mutant and wild-type plants we found only a small overlap of genes that are up-or downregulated after zeocin treatment (Figure 6a), which is in contrast to the high overlap after UV-C treatment (Figure 6b). Gene ontology term analyses of transcripts that are only responsive in the mutant and not the wild type or *vice versa* revealed that genes that are exclusively downregulated in the mutant are mainly involved in mitotic cell cycle regulation and microtubule movement (Figure 6c, d and Figure S3h). However, we also observed several regulators of the G2/M checkpoint control machinery to be affected by the zeocin treatment in the wild type, but not in the Δ*linda* mutant (Figure 6e). This could indicate that the inhibition of the cell cycle in the Δ*linda* deletion mutant is defective. Surprisingly, after UV-C treatment we found the opposite effect: while several G2/M checkpoint regulators were differentially expressed in the Δ*linda* mutant, they did not significantly respond in the wild type (Figure 6f). However, again we observed that the overall differences between the wild type and the mutant were rather low after UV-C treatment (Figure 6f). One striking observation is that *ATR* was strongly induced in the Δ*linda* mutant, while it was downregulated in the wild type during zeocin treatment (Figure 6e). A second observation was that the G2/M regulator *MYB3R3* was either not responsive or downregulated in the Δ*linda* mutant, which contrasts with the strong upregulation observed in the wild type in response to both UV-C and zeocin (Figure 6e, f).

**Figure 6.**
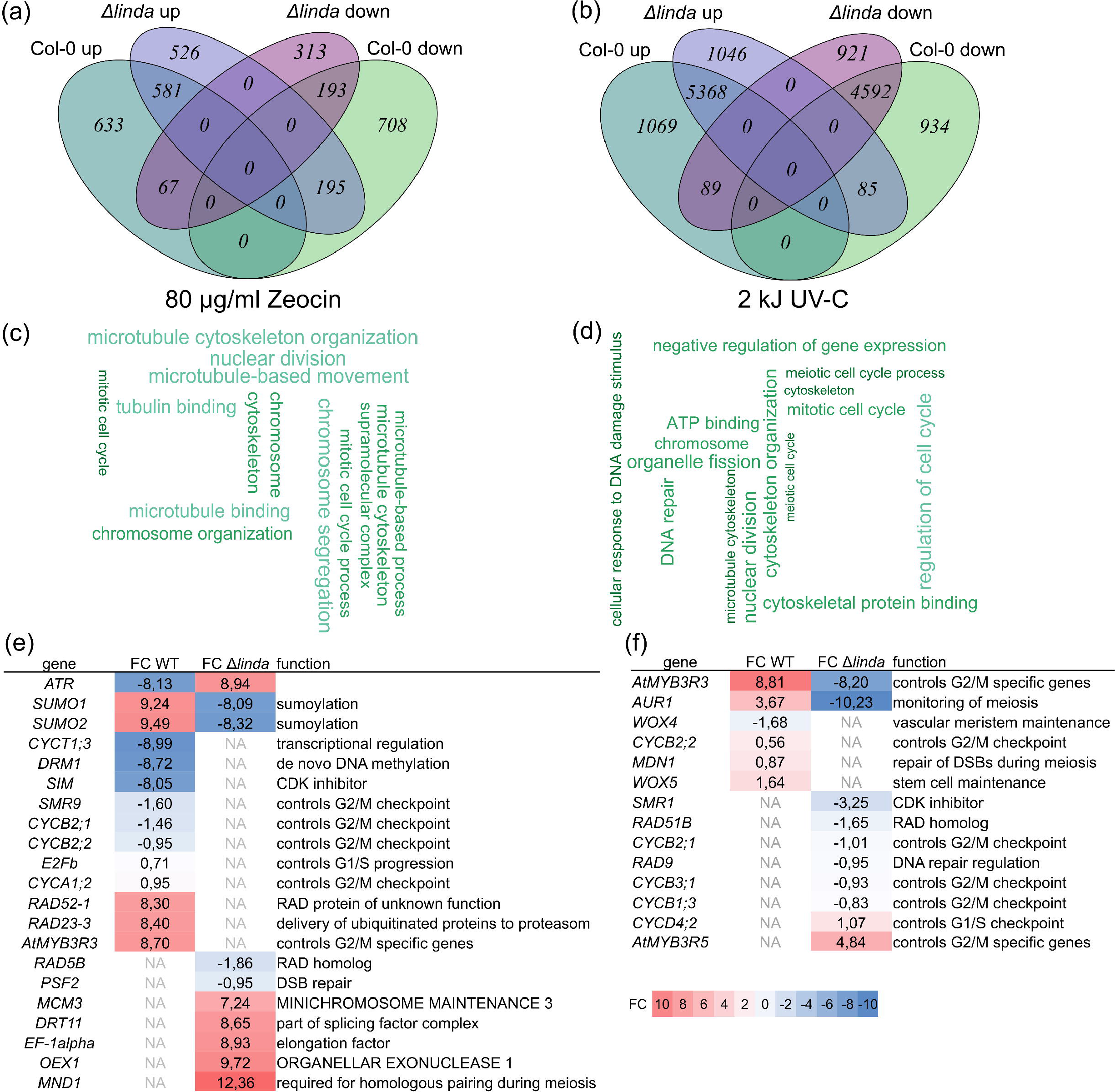
Differentially expressed genes in the wild type and the Δ*linda* mutants after DNA damage. (a-b) Overlap of up- and downregulated genes in wild-type and Δ*linda* mutant plants 3 h after treatment with zeocin (a) or UV-C (b). (c-d) Word cloud of gene ontology terms, downregulated in Δ*linda* but not in the wild type after zeocin (c) or UV-C treatment (d). (e-f) Identified putative target genes, differentially regulated in the Δ*linda* mutant versus the wild type, after treatment with zeocin (e) or UV-C (f). The measured fold change (FC) is indicated. NA, not available.

### The **Δ***linda* mutant is involved during the recovery from stem cell death

To further analyze the sensitivity of the Δ*linda* mutant towards different DNA-damaging stresses, we executed true leaf assays for both zeocin and UV-C treatment. Because the proportion of Δ*linda* seedlings developing true leaves after both treatments did not differ from the wild-type (Figure S4a, b), we investigated the root development of the Δ*linda* deletion mutant 24 h after treatment with high doses of zeocin. The Δ*linda* deletion mutant showed an increased area of cell death within the root meristem (Figure 7a, b), again suggesting a role for *LINDA* in the response to zeocin. We further examined if hyperactivation of cell death has an influence on the recovery of the roots following DNA damage, using the previously established root stem cell recovery assay that is based on a transient treatment for 24 h with 0.6 µg/ml bleomycin, followed by retransfer to recovery medium without bleomycin (Bisht *et al*., 2023). Using this setup, Δ*linda* mutants showed a reduced root growth in comparison to the wild-type (Figure 7c). Four days after the treatment, only 5% (1/20) of the roots were able to recover in the Δ*linda* mutant background, in contrast to 70% (14/20) roots in wild-type (Figure 4d, e). Thus, *LINDA* is an important factor for root recovery after treatment with DNA damaging agents.

**Figure 7.**
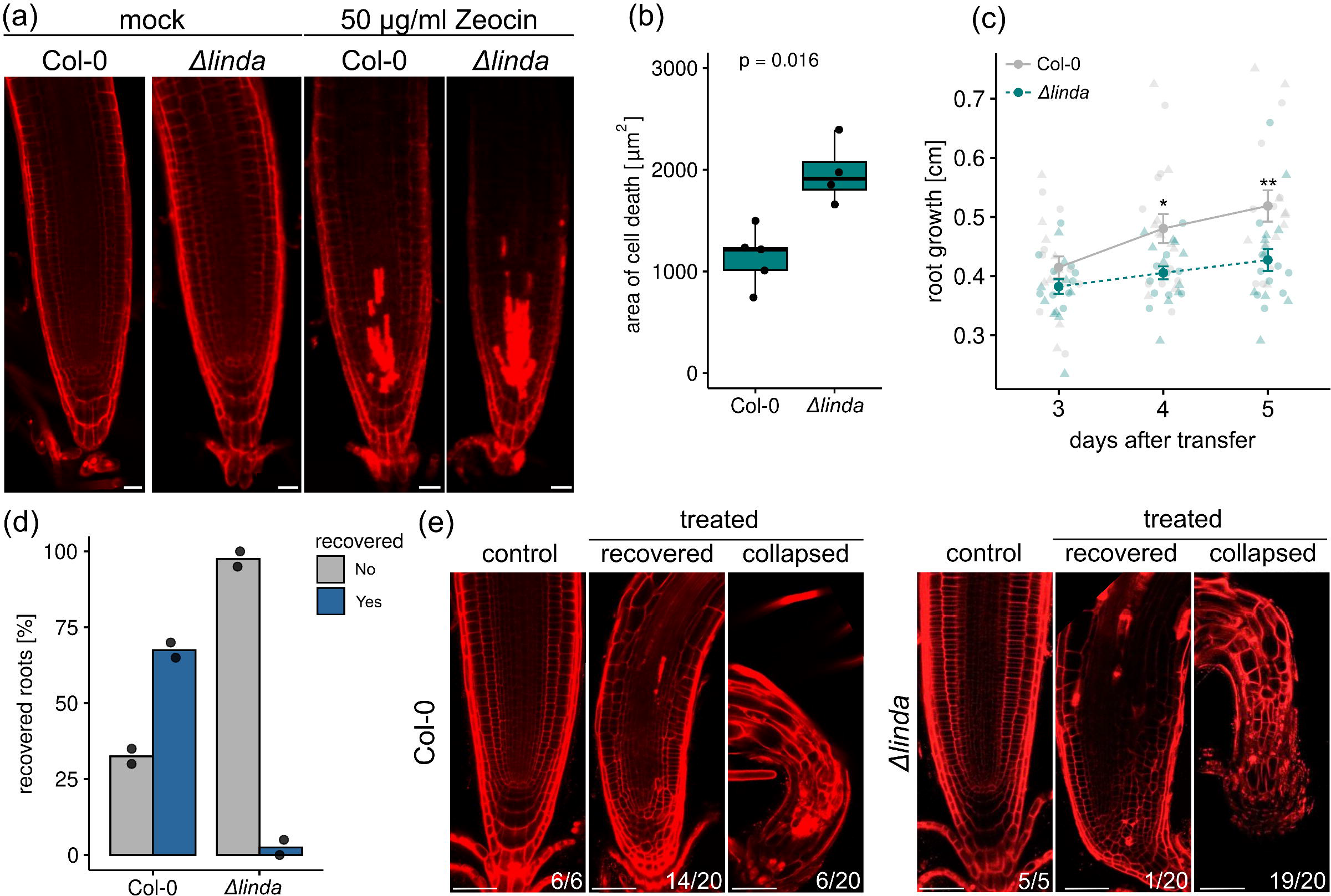
Analysis of the Δ*linda* deletion mutant. (a) Representative images of roots, stained with propidium iodide (PI). Seedlings, germinated on vertical MS plates without any additive, were transferred to vertical MS plates with or without 50 µg/ml zeocin for 24 h, seven days after germination. Scale bars = 50 µm. (b) Quantification of the area stained by PI, indicating dead cells. Statistical significance was evaluated by Wilcox Rank Sum Test (n = 4, p ≤ 0.05). (c) Root growth of seedlings that were transferred on bleomycin-containing medium for 24 h, five days after germination and subsequently transferred to medium without any additives. Root growth was measured at different days after retransfer to medium without any additives. Different shapes indicate for two repeats. Statistical significance was evaluated by Kruskal-Wallis Test, Bonferroni post-hoc (n ≥ 19, p ≤ 0.05 [*], p ≤ 0.01 [**]). (d) The percentages of recovered (yes) and collapsed (no) roots from two biological repeats were determined four days after retransfer to bleomycin free medium. (e) Representative images and numbers of recovered and collapsed roots after staining of the roots with PI. The total number of monitored roots is also indicated. Scale bars = 50 µm.

## DISCUSSION

### *LINDA* is an early DNA double-strand break-induced lncRNA

In humans, already 13 lncRNAs have been identified with a function in maintaining genome integrity and stability (Li *et al*., 2021), whereas in plants no lncRNAs acting in these processes are reported yet. We here describe the characterization of the first lncRNA with a role in fine-tuning the DDR in the model plant *A. thaliana*.

We identified the lncRNA *LINDA* by meta-analysis of bulk RNA-Seq results. We found *LINDA* to be strongly induced by different stresses inducing primarily DSBs, including zeocin and bleomycin, by UV-C, which causes predominantly the formation of pyrimidine-dimers, as well as by X-rays (Wang *et al*., 2016) and gamma-rays (Bourbousse *et al*., 2018). The pyrimidine dimers caused by UV-C can be either directly reversed by photoreactivation (Jiang *et al*., 1997) or removed by nucleotide excision repair (NER). As the NER mechanism includes the formation of DNA single- and double-strand breaks to remove the damaged bases (Ries *et al*., 2000, Molinier *et al*., 2004), the induction of *LINDA* after UV-C irradiation could eventually also be caused by the formation of DSBs. Differently, the applied UV-B doses might not have been sufficient to produce DSBs and thus did not trigger *LINDA* induction. Contrary to DSB-inducing agents, *LINDA* does not respond to mitomycin C, which causes DNA cross-links and thereby replication stress. Consequently, we hypothesize that *LINDA* responds specifically to DNA DSBs.

Based on the expression kinetics, *LINDA* can be attributed to the early-induced gene cluster proposed by Bourbousse *et al*. (2018). This cluster contains mainly genes that specifically respond to genotoxic stress and are involved in DNA repair, DNA metabolism, gene regulation, and cell cycle control, like *BRCA1*, *SIAMESE-RELATED 5 (SMR5)* and *SMR7*. Moreover, the *LINDA*-flanking and co-regulated gene *At3g45730* was also found within this particular cluster (Bourbousse *et al*., 2018). Although coexpression does not imply functional relevance, it is an important hint towards a functional relationship between *LINDA*, the hypothetical At3g45730 protein and the DDR pathway. Another important indication towards a function of *LINDA* in the plant DDR is the direct regulation of *LINDA* transcription by SOG1 in an ATM-dependent manner. ChIP-seq experiments with FLAG-tagged SOG1 resulted in 310 potential target genes for the SOG1 transcription factor, including *At3g45730* (Bourbousse *et al*., 2018). The lncRNA *LINDA* was not annotated as a putative SOG1 target in the screen from Bourbousse *et al*. (2018), however based on our findings, we can add *LINDA* to the list of SOG1 target genes. Among those potential SOG1 target genes, the majority is involved in DDR-related mechanisms. We therefore hypothesize that *LINDA*, being a SOG1-target gene, is also involved in the plant DDR.

We showed that in *atm* and *sog1-1* mutant plants, the response of *LINDA* to zeocin is lost. In contrast, in these mutants, *LINDA* is still induced by UV-C irradiation, although the induction is significantly decreased relative to wild-type plants. UV-C can produce a multitude of different DNA lesions, and the repair of these lesions presumably involves both ATR and ATM. On the other hand, zeocin does primarily lead to the formation of DSBs, which are preferentially regulated by ATM (Culligan *et al*., 2006). We speculate that *LINDA* might be regulated by other transcription factors, besides SOG1, in response to UV-C radiation. As it was recently shown that target genes of the E2FA/E2FB transcription factors overlap with those of SOG1 (Nisa *et al*., 2023). *LINDA* might be additionally controlled by E2F transcription factors, at least after UV-C treatment.

### *LINDA* regulates its putative target gene in *cis*

Many intergenic lncRNAs regulate the gene expression of their target genes in *cis* (Kopp and Mendell, 2018, Rai *et al*., 2019, Lucero *et al*., 2021). We have unraveled a strong connection between *LINDA* and the 644-bp downstream located, flanking gene *At3g45730*. Both genes are induced in response to various DNA damaging stresses, and both are regulated by the ATM-SOG1 pathway. Moreover, the Δ*linda* mutant exhibits a hyperactivation of *At3g45730*, thus underlining the putative *cis* regulatory impact of the lncRNA towards its flanking gene. However, it became evident that *LINDA* might be a conditional regulator of its putative target gene, because *At3g45730* is still expressed at wild-type levels in the Δ*linda* mutant after UV-C treatment.

We assume that *LINDA* could either regulate its flanking gene in a dose-dependent manner, has a different set of putative target genes after UV-C treatment or has another, yet unidentified, intermediate factor. The first hypothesis is supported by the observed dose dependency of the *LINDA* gene expression. On the other hand, the second hypothesis is strengthened by the transcriptome analyses, were only a minor overlap between differentially regulated genes was found for the UV-C-versus the zeocin-treated Δ*linda* samples. Lastly, the partial induction of *LINDA* after UV-C treatment, as well as the hyperactivation of *LINDA* in the *atr* mutant background could argue for the third hypothesis.

Interestingly, both *LINDA* and *At3g45730* share high sequence similarity due to a small putative ORF within the *LINDA* sequence. We hypothesize that *LINDA* originated from *At3g45730* by gene duplication and subsequently underwent pseudogenization. A common feature of pseudogenization is the remnant of a putative ORF (Kapusta and Feschotte, 2014), like in the case of *LINDA*. This remnant ORF makes it difficult to estimate if the gene is a true lncRNA or not (Kapusta and Feschotte, 2014). Intriguingly, we found this gene duplication and putative pseudogenization also in several other Brassicales and Malvales species. However, whether those putative gene duplicates are indeed functional lncRNAs remains to be confirmed in the future. To date, profound knowledge about the evolution of lncRNAs is lacking and it is only assumed that gene duplication can give rise to new lncRNAs but this process does probably not contribute significantly to the origin of most lncRNAs (Kapusta and Feschotte, 2014). However, studies about the origin of lncRNAs mainly focus on the evolution of mammalian lncRNAs (Elisaphenko *et al*., 2008, Kapusta and Feschotte, 2014, Marques and Ponting, 2014, Hezroni *et al*., 2017, Waters *et al*., 2021). Therefore, it cannot be excluded that plants have their own evolutionary trails for lncRNAs.

Based on the high sequence similarity between both genes, it is tempting to speculate that *LINDA* might function as decoy or target mimicry for activators of *At3g45730* and consequently might reduce a potentially unwanted overaccumulation of the hypothetical protein. LncRNAs functioning as target mimicry have been described before, like for example in the case of *IPS1* that sequesters miRNA399 by target mimicry (Franco-Zorrilla *et al*., 2007, Jabnoune *et al*., 2013). This regulation could also be mediated via RNA:DNA triplex formation, which has been shown to be important for activation and repression of the target genes, at least in mammals (Schmitz *et al*., 2010, O’Leary *et al*., 2015).

### *At3g45730* as potential regulator of cell death

We showed that *At3g45730* is, similar to *LINDA*, highly induced by various DNA-damaging stresses and has been identified in multiple screens focusing on ATM (Culligan *et al*., 2006) or SOG1 (Bourbousse *et al*., 2018) target genes, thus underlining its potential role in the DDR. However, *At3g45730* was also found in two screens to be upregulated by antimycin A and oligomycin (Van Aken and Pogson, 2017, Shapiguzov *et al*., 2019). Both antimycin A and oligomycin trigger mitochondria dysfunction, which can ultimately lead to the formation of ROS and the induction of cell death, as mitochondria are the drivers for programmed cell death (Yao *et al*., 2004). Moreover, bleomycin triggers the fragmentation of mitochondrial DNA, accompanied by ROS burst and cell death, at least in mammals (Yeung *et al*., 2015, Suryadevara *et al*., 2019). It is thus conceivable that the hypothetical At3g45730 protein is linked to mitochondrially derived ROS or programmed cell death signals. A well-known example of a peptide functioning in programmed cell death is the 25-amino acid-long kiss of death (KOD) peptide that might be linked to cellular ROS levels and is involved in the depolarization of the mitochondrial membrane (Blanvillain *et al*., 2011). Although a loss-of-function line would help to study the molecular role of At3g45730, to date, no transgenic lines are available and we failed to create CRISPR/Cas lines, which might indicate that a *knockout* of *At3g45730* is lethal.

### *LINDA* is involved in the proper response to high doses of genotoxic stress

Phenotypical analysis of the Δ*linda* mutant yielded in an increased area of cell death in response to high doses of zeocin. The increased cell death is most likely one of the reasons for the strong reduction of the recovery rate of Δ*linda* seedlings after destruction of the root meristem by high doses of bleomycin. Moreover, genome wide transcription studies indicated that zeocin causes a more severe transcriptome rearrangement in the Δ*linda* deletion mutant than UV-C. After zeocin treatment, the Δ*linda* mutant showed defects in the transcriptional activation/repression of several cell cycle regulators. In contrast, the relative number of deregulated genes after UV-C treatment was low. Moreover, we could not find an enrichment for genes linked to cell death in the Δ*linda* mutant. Further studies must prove if *LINDA* is directly involved in the cell cycle regulation, or if this is only the consequence of an affected DNA repair machinery and/or DDR signaling and how cell death is linked to these mechanisms. One important finding in this regard was that the *LINDA* deletion led to the hyperactivation of *ATR* after zeocin treatment, which might be connected to the hyperactivation of *LINDA* in the *atr* mutant. Thus, *LINDA* might be needed for the accurate performance/activation/signaling of the DDR pathway in response to different DNA-damaging stresses.

In conclusion, we could show the regulation of *LINDA* by ATM and SOG1 in response to DNA damage as well as the importance of *LINDA* in the DDR machinery. The exact molecular function of *LINDA* in the recognition and repair of damaged DNA will have to be investigated in future studies.

## MATERIALS AND METHODS

### Plant material and growth conditions

*A. thaliana* seeds were sterilized with 70% ethanol for 2 min, washed with water twice, and sown on half-strength Murashige & Skoog (MS) medium (½ MS, 100 mM MES, pH 6.5) containing 0.8% (w/v) and 1% (w/v) plant culture agar for horizontal and vertical growth plates, respectively. Afterward, plants were grown under short-day conditions (10-h light/ 14-d darkness, 100 µE, 20°C, 60% humidity). The ecotype Columbia-0 (Col-0) was used as the wild type for all analyses. The *sog1-1*, *atm-2* and *atr-2* mutants were described before (Garcia *et al*., 2003, Preuss and Britt, 2003, Culligan *et al*., 2004). The *atr-2* mutant (SALK_032841) seeds were kindly gifted by Roman Ulm and Richard Chappuis (University of Geneva, Geneva, CH).

### Plant treatments

To determine the gene expression of wild-type and mutant *A. thaliana* plants, 21-day-old seedlings were transferred to ½ liquid MS medium (½ MS, 100 mM MES, pH 6.5) with or without 1 µg/ml bleomycin, 10 µg/ml mitomycin-C, 40 µg/ml, or 80 µg/ml zeocin. Samples were harvested after 90 min, 3 h, 6 h, or 24 h of incubation. In the case of UV radiation, the seedlings grown on plates were directly treated with 2 kJ of either UV-A, UV-B, or UV-C. The dose was measured with UVTOUCH (sglux). Afterward, the plants were transferred back to short-day conditions. Samples were harvested at 30 min, 60 min, 90 min 3 h or 6 h after the treatment.

### Construction of **Δ***linda* deletion mutants by CRISPR/Cas12a

The vectors to construct the CRISPR/Cas12a deletion mutants were kindly provided by P. Schindele and H. Puchta from the Karlsruher Institut für Technologie. We used the dual guideRNA (gRNA) approach to target either the transcribed region of *LINDA* or the intergenic sequences downstream of the transcribed region. The gRNA oligonucleotides (Table S1) were synthesized by Eurogentec. Both gRNAs were introduced into the entry vector pEn-RZ_Lb-Chimera, which was digested prior with BbsI (NEB). The first gRNA was inserted into the destination vector pDe-EC-ttLbCas12a by the LR-clonase (Thermo Fisher). The second gRNA was amplified from the entry vector with PS1 and PS2 primers (Table S1). Next, the pDe-EC-ttLbCas12a vector, containing the first gRNA, was linearized with BamHI. The final vector was constructed by Gibson Assembly (NEB Hifi DNA Assembly Mastermix) using the linearized pDe-EC-ttLbCas12a vector and the amplified second gRNA. The final vector was transformed into *A. thaliana* with Agrobacteria using the floral-dip method. Positive transformants were selected on MS plates containing the selection marker gentamicin (60 µg/ml). The presence of the T-DNA was confirmed by PCR using one pDe-EC-ttLbCas12a vector-specific primer and on primer specific for the second gRNA (Table S1). The seeds of each positive transformed line were collected separately. In the next generation, the number of T-DNA insertions was determined by growing plants again on MS plates containing gentamicin (60 µg/ml). Lines with a survival rate of 75% on the gentamicin-containing plates were propagated on MS plates without a selection marker. Next, the deletion of the *LINDA* gene was characterized using *LINDA*-specific primers, while the absence of the T-DNA was confirmed as described before. Plants with a partial deletion of *LINDA* and without the T-DNA were selected for further analysis. To confirm the absence of the T-DNA in the selected lines do not comprise the T-DNA anymore, seeds were again grown on gentamicin-containing MS plates.

### True leaf assay

True leaf assays were performed as previously described for zeocin (Ryu *et al*., 2019) and UV-C (Rosa and Mittelsten Scheid, 2014).

### Confocal imaging and determination of cell death area

Seedlings were germinated on ½ MS medium without any additives for seven days and subsequently transferred to ½ MS medium containing 50 µg/ml zeocin. After 24 h, the seedlings were stained for 3 min with propidium iodide (5 µg /ml PI) and the root meristem was imaged by confocal laser-scan microscopy (Zeiss 710; λ_ex_ = 488 nm; λ_em_ = 600 – 650 nm). The area of dead cells, stained by the penetrated PI, was measured for at least four independent roots and quantified with ImageJ (version 1.53t).

### Recovery assay

*A. thaliana* seeds were sown on horizontal ½ MS plates and vernalized for two days at 4°C. Seedlings were grown for four days under standard short-day conditions. Seedlings with equal root lengths were treated for 24 h on ½ MS medium, containing 0.6 µg/ml bleomycin, and transferred to medium without bleomycin. The root growth was tracked for five consecutive days, and the root apical meristem was visualized by confocal laser-scan microscopy (Zeiss 710; λ_ex_ = 488 nm; λ_em_ = 600 – 650 nm) after staining with PI (5 µg/ml) for 3 min.

### RNA isolation and cDNA synthesis

The used RNA isolation protocol was performed as previously described (Oñate-Sánchez and Vicente-Carbajosa, 2008). To increase solubility, after resuspension of the isolated RNA in DEPC-treated water, RNA was frozen for 1 min in liquid nitrogen and subsequently incubated for 10 min at 65°C. Next, 7.5 µg RNA were treated with 1 U DNase for 30 min at 37°C. The reaction was stopped by adding 1 µl 25 mM EDTA and heating the samples (10 min, 65°C). Next, the RNA was diluted with 80 µl DEPC-treated water and precipitated by adding 10 µl 3 M sodium acetate and 400 µl 96% ethanol. To increase yield, the RNA was incubated for 20 min at −20°C, followed by centrifugation for 20 min at 16.000 g and 4°C. The pellet was washed with 500 µl of 70% ethanol and air-dried. The RNA was resuspended in 15 µl DEPC-treated water. The samples were frozen in liquid nitrogen and subsequently incubated for 10 min at 65 °C. Next, 500 ng of RNA were used for cDNA synthesis according to the manufacturer’s instructions (roboklon, smaRT).

### RT-qPCR

RT-qPCR reaction mixtures were set up according to the manufacturer’s instructions (Blue ŚGreen, Biozym) and reactions were carried out on a CFX96^TM^ Touch thermocycler (Bio-Rad). As a control for contamination with genomic DNA, two primers specific for the AGAMOUS-LIKE 68 intron (AGL68) were used (Table S1). Only samples with a cycle threshold (Ct) value for AGL68 > 30 were used for further analysis. Moreover, by using two primer pairs, which are specific for either the 5’ or the 3’ end of the glyceraldehyde-3-phosphate-dehydrogenase (GAPDH), the integrity of the cDNA was determined (Table S1). If the difference in Ct values of GAPDH5’ and GAPDH3’ was above 1.5, samples were excluded from further analysis. UBIQUITIN 10 (UBI) was used as a reference gene for normalization (Table S1).

### RNA-Seq analyses

For the RNA-Seq analyses, seedlings were grown for two weeks under standard light conditions and either treated with 80 µg/ml zeocin as described above in ½ liquid MS, or subjected to 2 kJ of UV-C. Three hours after treatment, the RNA was isolated using the Quick-RNA Plant Kit (Zymo research). One µg of isolated RNA was sent to Macrogen for sequencing. The resulting sequences were analyzed with the Galaxy platform, following the workflow provided by Dr. Lieven Sterck (RNA-seq Paired End Reads, Salmon v20211020, The Galaxy Community, 2022) The acquired gene levels for each sample were then used for the analysis of differentially expressed genes, using the DEseq2 package from Bioconductor (Love *et al*., 2014).

### Meta-analysis

The following datasets, used for meta-analysis, were obtained from Gene Expression Omnibus (GEO, https://www.ncbi.nlm.nih.gov/geo/): GSE111062 - 1200 µE HL (Huang *et al*., 2019), GSE143762 - 0.1 kJ UV-C (Czarnocka *et al*., 2020), GSE158992-28 °C (Lee *et al*., 2021). Additionally, the RNA-Seq dataset from Wang *et al*. (2016) was included. The analyses of the acquired datasets were performed with the DESeq2 package from Bioconductor (Love *et al*., 2014). Significantly regulated genes were selected (p < 0.05). The gene model annotations for the significantly regulated genes were obtained from Araport_11. The following terms were considered as lncRNA: other RNA, antisense lncRNA, novel transcribed regions, and lncRNA. Venn diagrams were generated with the VennDiagram package (version 1.7.1) from The Comprehensive R Archive Network (CRAN).

### Alignment

Alignments were constructed with ClustalW by the Molecular Evolutionary Genetics Analysis (MEGA-X) software (version 10.0.5). The phylogenetic tree was constructed with the maximum likelihood method by MEGA-X. Alignments were visualized with Jalview (version 2.11.1.7).

### Phylogenetic analysis

Similar sequences were identified by the Standard Nucleotide Basic Local Alignment Search Tool (BLASTN), using either the transcribed sequences of *At3g00800* or *At3g45730* as the query. The intergenic regions of the identified putative homologs were obtained from the National Center for Biotechnology Information Genome Data Viewer. The following genome sequence assembly data were used: *Arabidopsis lyrata* (v.1.0), *Arabidopsis thaliana* (TAIR10.1), *Brassica napus* (Bra_napus_v2.0), *Brassica rapa* (CAAS_Brap_v3.01), *Camelina sativa* (Cs), *Capsella rubella* (Caprub1_0), *Eutrema salsugineum* (Eutsalg1_0), *Gossypium arboreum* (Gossypium_arboreum_v1.0), *Gossypium hirsutum* (Gossypium_hirsutum_v2.1), *Tarenaya hassleriana* (ASM46358v1), *Theobroma cacao* (Criollo_cocoa_genome_V2). We searched for SOG1 binding motifs within the selected intergenic regions with SnapGene Viewer (version 5.3.2).

### Statistical analysis

All statistical analysis were performed with the RStudio Server (Version 1.3.1073 RStudio Team. (2015). RStudio: Integrated Development Environment for R. Boston, MA. Retrieved from http://www.rstudio.com/). Assumptions for normality and homogeneity of variance were tested by the Shapiro and Levene Test, respectively. Depending of the outcome of the assumption analysis, either an analysis of variance (ANOVA), Kruskal-Wallis, or Student’s *t*-test were performed. Bonferroni correction was used for p-value adjustment. The data visualization was done with the RStudio Server, using the ggplot2 (Wickham, 2016) and ggpubr packages (Kassambara, 2023).

## ACCESSION NUMBERS

AT3G45730; AGL68: AT5G65080; AGO2: AT1G31280; ATM: AT3G48190; ATR: AT5G40820; BRCA1: AT4G21070; GAPDH: AT3G26650; RAD51: AT5G20850; SOG1: AT1G25580; LINDA: AT3G00800; UBI10: AT4G05320

## ACKNOWLEDGEMENTS

We thank Dr. Patrick Schindele and Prof. Dr. Holger Puchta from the Karlsruher Institut für Technologie for providing the used vectors for construction of the CRISPR/Cas12 mutants, and Richard Chappuis and Prof. Dr. Roman Ulm from the University of Geneva for providing the used *atr-2* seeds. We thank Dr. Valentin Hammoudi and Asst. Prof. Dr. Andrew Nelson for valuable discussions throughout the project and during manuscript preparation. We thank Dr. Annick Bleys for her help in improving the manuscript.

## SUPPORTING INFORMATION

Additional Supporting Information may be found in the online version of this article.

**Figure S1: The percentage of all transcripts and identified lncRNAs in the performed meta-analysis that are responsive to only one treatment**

**Figure S2: Sequence alignment of LINDA and At3g45730.**

**Figure S3: Transcriptome analyses of the wild-type and Δlinda mutant response to DSBs.**

**Figure S4: True leaf assay Table S1: Used primers**

