## Supplemental data for "The long non-coding RNA *LINDA* restrains cellular collapse following DNA damage in *Arabidopsis thaliana*"

**SUPPORTING INFORMATION**

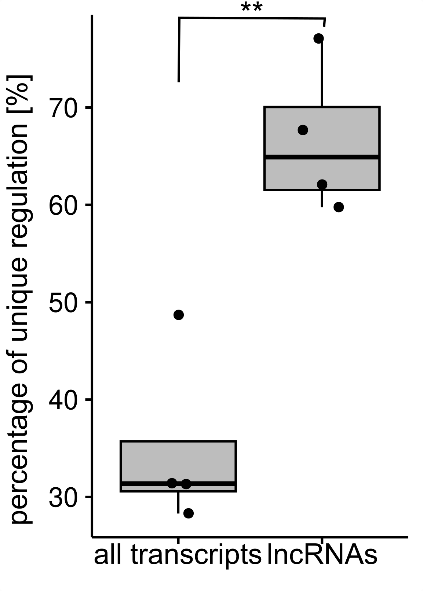

**Figure S1. The percentage of all transcripts and identified lncRNAs in the performed meta-analysis that are responsive to only one treatment (Student’s *t*-test, p < 0.01 [**]).**

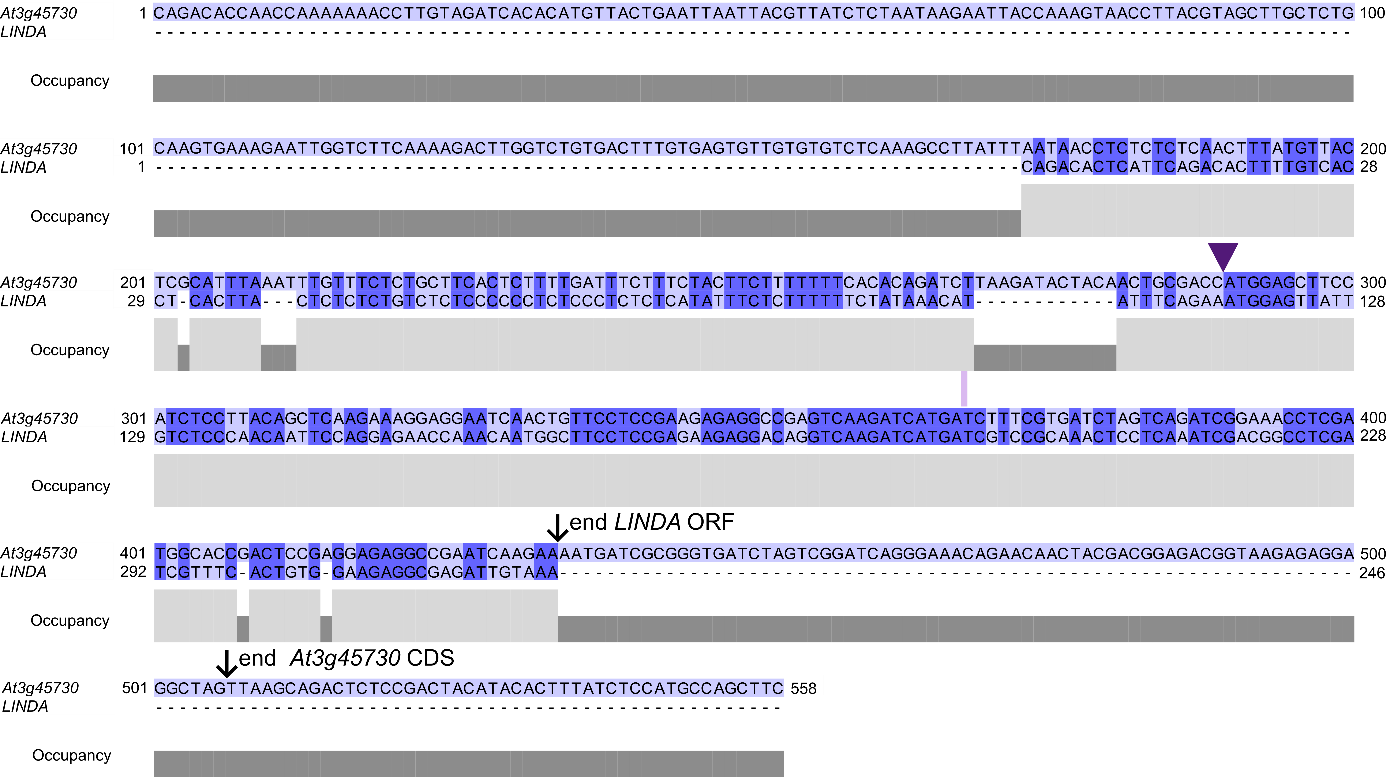

**Figure S2. Sequence alignment of *LINDA* and *At3g45730.*** Identical nucleotides are highlighted in dark blue. The triangle marks the start of the coding sequence of *At3g45730* and of the short open reading frame (ORF) of *LINDA*. Black arrows indicate to the end of the coding sequence and the ORF.

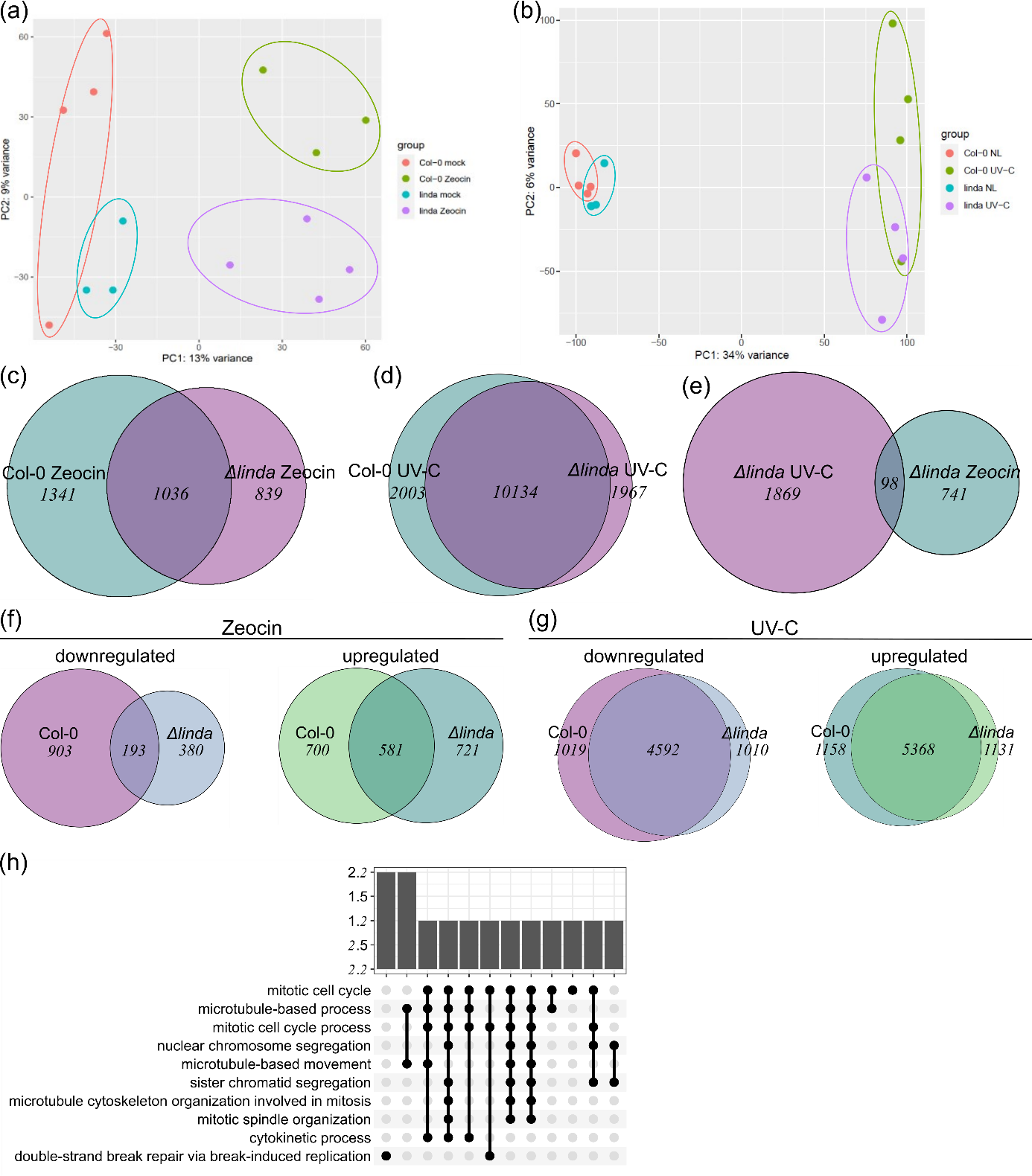

**Figure S3. Transcriptome analyses of the wild-type and *Δlinda* mutant response to DSBs.** (a-b) Principle component analyses (PCA) of the wild type (Col-0) and *Δlinda* transcriptomes after treatment with or without (mock) 80 µg/ml zeocin (a) or no light (NL) or 2 kJ UV-C (b). (c-d) Overlap of significantly (p ≤ 0.05) differentially expressed genes in the *Δlinda* mutant and wild type after zeocin (c) or UV-C treatment (d). (e) Overlap of genes that were only significantly (p ≤ 0.05) deregulated in the *Δlinda* deletion mutant. (f-g) Venn diagram of significantly (p ≤ 0.05) up- or downregulated genes in the *Δlinda* mutant versus the wild type after zeocin (f) or UV-C treatment (g). (h) Upsetplot of genes deregulated in the *Δlinda* mutants but not in the wild type after zeocin treatment.

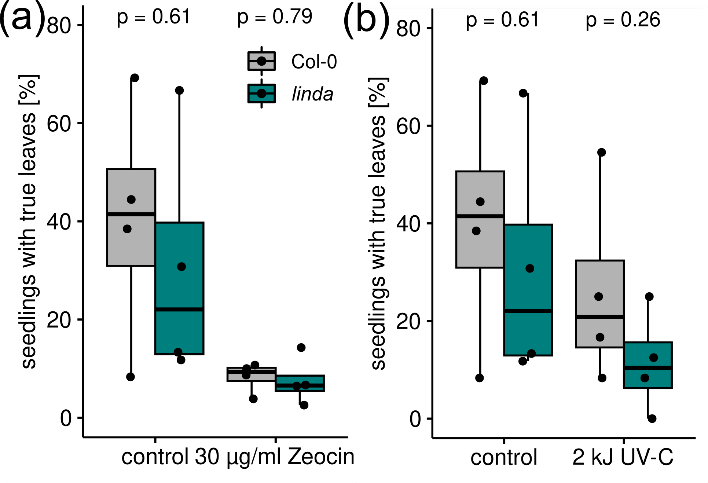

**Figure S4. True leaf assay.** (a-b) Percentage of *Δlinda* mutant seedlings developing true leaves after germination on medium containing 30 µg/ml zeocin (a) or after treatment with 2 kJ UV-C (b). Statistical significances were evaluated by Student’s *t*-test (n = 4).

**Table S1. Overview of the primers used for this study**

| name | 5’ – 3’ sequence | purpose |
| --- | --- | --- |
| UBI10 fwd | GGCCTTGTATAATCCCTGATGAATAAG | RT-qPCR |
| UBI10 rev | AAAGAGATAACAGGAACGGAAACATAGT | RT-qPCR |
| GAPDH5’ fwd | TTGGTGACAACAGGTCAAGCA | RT-qPCR |
| GAPDH5’ rev | AAACTTGTCGCTCAATGCAATC | RT-qPCR |
| GAPDH3’ fwd | TCTCGATCTCAATTTCGCAAAA | RT-qPCR |
| GAPDH3’ rev | CGAAACCGTTGATTCCGATTC | RT-qPCR |
| AGL68 fwd | TTTTTTGCCCCCTTCGAATC | RT-qPCR |
| AGL68 rev | ATCTTCCGCCACCACATTGTAC | RT-qPCR |
| LINDA fwd | TTCAGAAATGGAGTTATTGTCTCC | RT-qPCR |
| LINDA rev | GATTTGAGGAGTTTGCGGAC | RT-qPCR |
| At3g45730 fwd | TAAGATACTACAACTGCGACCA | RT-qPCR |
| At3g45730 rev | GTCTGCTTAACTAGCCTCCT | RT-qPCR |
| BRCA1 fwd | ACCAGAATGCAGATGGGACAATG | RT-qPCR |
| BRCA1 rev | TAGGCTGAGAGTGCAGTGGTTC | RT-qPCR |
| RAD51 fwd | GCGCAAGTAGATGGTTCAGC | RT-qPCR |
| RAD51 rev | TTCCTCAACGCCAACCTTGT | RT-qPCR |
| PS1 | ccggaccgactagtggatccAAAGCAGGCTACGCGTCCT | cloning |
| PS2 | aggtcgactctagaggatccAAAGCTGGGTCCTCAGGAC | cloning |
| LINDA CRISPR fwd | CGTATAACCATGCCCTGACC | genotyping |
| LINDA CRISPR rev | GTTAGGCTTTATGCATGTCCC | genotyping |
| LINDA guide 2 fwd | AGATAGAAATGGAGTTATTGTCTCCCAA | cloning |
| LINDA guide 2 rev | GGCCTTGGGAGACAATAACTCCATTTCT | cloning |
| LINDA guide 4 fwd | AGATATACGTGATACATGAGATTACATC | cloning |
| LINDA guide 4 rev | GGCCGATGTAATCTCATGTATCACGTAT | cloning |
